# Spatial Transcriptomics Reveals that the Local Immune Response to Placental Guinea Pig Cytomegalovirus Infection is Driven by Chemokine Signaling

**DOI:** 10.1101/2025.08.01.668085

**Authors:** Zachary W. Berkebile, Colleen L. Forster, Emma Stanley, Fernanda M. Rodriguez, Adam C. Herman, Micah D. Gearhart, Craig J. Bierle

## Abstract

Cytomegalovirus infection can disrupt placental development and function either by directly infecting placental cells or by eliciting a pathogenic immune response. The relative contributions of these two mechanisms to adverse pregnancy outcomes remains poorly understood. In this study, we used spatial transcriptomics to quantify host and viral gene expression at the maternal-fetal interface at near single-cell resolution. Guinea pig cytomegalovirus (GPCMV) infection after mid-gestation causes focal infections at the base of the main placenta. Samples for spatial transcriptomics were collected from guinea pigs infected with GPCMV at 35 days gestation. Viral loads and the location of infected cells in placentas were assessed using virus-specific droplet digital PCR and *in situ* hybridization at 21 days post-infection. Representative placentas were sectioned onto Visium Spatial Gene Expression Slides and sequencing libraries were prepared from six infected and six uninfected tissue sections. Spatial transcriptomes from 33,687 55-µm spots were generated and used in subsequent analyses. To assess how infection affected gene expression at the maternal-fetal interface, a combination of graph-based clustering and manual classification was used to assign spatial transcriptomes to clusters representative of different anatomic regions. Infection dysregulated more transcripts in the decidua and junctional zone than in the labyrinth or non-capillarized syncytium. Notably, infection downregulated transcripts involved in lipid metabolism and upregulated transcripts involved in antiviral defense and chemokine signaling. A second analysis compared the spatial transcriptomes of GPCMV-infected cells and their immediate microenvironment with similar regions in uninfected placentas. A transcriptional signature indicative of immune activation was clear in this comparison, and the local placental response to cytomegalovirus infection was driven by upregulated chemokine signaling. Thus, spatial transcriptomics revealed regional patterns of gene expression in the guinea pig placenta and illuminated how the host response to GPCMV may compromise placental function.

**Author Summary:** The placenta supports fetal development while also acting as an immune barrier against bloodborne pathogens. Cytomegalovirus (CMV), the most common viral cause of congenital infections and preventable neurologic disability in children, evades host defenses to infect the placenta. How CMV affects the placenta and fetal health remains poorly understood. Using a guinea pig model of CMV infection during pregnancy and spatial transcriptomics, a recently-developed method that enables gene expression to be studied at near-cellular resolution, this study compared normal and infected placentas. The effects of infection on directly infected cells and their immediate environment and indirect effects that occur at more distant sites were revealed. This information may inform the development of therapies to improve placental function after CMV infection.

## Introduction

Human cytomegalovirus (HCMV) is a ubiquitous herpesvirus and the most frequent cause of congenital viral infections [1]. Congenital cytomegalovirus (cCMV) infections occur in roughly 1 in 200 pregnancies and approximately 8000 children are disabled by cCMV each year in the United States [2]. Placental infection plays a central role in the pathogenesis of cCMV. The placenta is critical for supporting fetal development, facilitates the development and maintenance of maternal tolerance, and acts as an immunologic barrier protecting the developing offspring from bloodborne pathogens. A myriad of antiviral defenses, including the constitutive expression of pattern recognition receptors and type III interferon, prevent most viruses from replicating in the placenta and/or infecting the fetus [3]. HCMV can infect both the maternal decidua and the placenta, placental infection likely precedes congenital infection, and some adverse pregnancy outcomes, including fetal growth restriction and stillbirth, can be caused by placental HCMV infections that do not transmit to the fetus [4–7].

How HCMV infection affects placental development and function remains poorly understood [8]. In clinical samples, HCMV antigens are most often detected in the decidua, in cytotrophoblasts located in floating villi and cell columns, and in the villous stroma and fetal capillaries [9, 10]. Villous malformations have been noted in cases of early fetal demise after congenital HCMV infection, and it has been proposed that infection can interfere with trophoblast proliferation and differentiation to compromise the early placental development [4]. Other less severe pathologies, including a hypoxia-like syndrome, are more frequent and observed when placental infection occurs later in pregnancy [5, 11]. Isolated primary extravillous trophoblasts, cytotrophoblasts, and trophoblast progenitor cells can be productively infected by HCMV *in vitro* [12–15]. Infection reduces the capacity of cytotrophoblasts to proliferate, limits the differentiation of trophoblast progenitor cells, and interferes with the anchoring of villous explants *in vitro* [16, 17]. HCMV can infect trophoblast stem cells, where the virus dysregulates the expression of transcripts involved in differentiation, but these infections are not productive, have no apparent effect on the proliferation of the trophoblast stem cells, and viral genomes are not maintained after passage [18, 19]. Similar observations have been made in trophoblast organoids, which also do not support productive HCMV replication [20].

The inflammatory response to placental infection can cause pregnancy loss, growth restriction, neurocognitive impairment, and developmental delay [21]. Proinflammatory cytokines accumulate in amniotic fluid, the placenta, and maternal sera during congenital HCMV infections [22–24]. Similar inflammatory responses occur when tissue explants and organoids are infected with HCMV *in vitro* [20, 25, 26]. Therapeutic or genetic interference with inflammatory signaling pathways can rescue murine pregnancies after infection, exposure to pathogen associated molecular patterns, or cytokines treatments [27, 28]. The species-specificity of HCMV and inability of mouse cytomegalovirus to cause placental and congenital infections has limited our ability to experimentally study whether the inflammatory response to placental infection is a cause of fetal harm [1, 29].

Guinea pigs are the most widely used animal model of cCMV infection. Guinea pigs have ∼65-day gestational periods and highly-developed, precocious offspring [30]. Major guinea pig organ and immune systems develop *in utero* to an extent that is more like humans than the mouse or rat. Guinea pig cytomegalovirus (GPCMV, also known as Caviid herpesvirus 2) was isolated from the salivary gland of an infected guinea pig and primary GPCMV infection during pregnancy leads to the sequential infection of the placenta and fetus [31–35]. Fetal outcomes are affected by the timing of infection during pregnancy [36, 37]. Notably, maternal infection after mid-gestation causes high rates of stillbirth and fetal growth restriction, and lesions, including ischemic injury and areas of necrosis, are observed in GPCMV-infected placenta at 14 days post-infection (dpi) and later [33]. We recently compared how maternal GPCMV infection either late in the embryonic period (21 days gestation [dGA]) or after mid-gestation (35 dGA) affected the placenta [38]. While there were no significant differences in viral genome abundance between these two groups at 21 dpi, bulk RNA-seq revealed that significant changes in placental gene expression were largely restricted to the animals that had been infected at 35 dGA. *In situ* hybridization further revealed differences in the location of infected cells in older and younger placenta. These observations led us to hypothesize that the guinea pig placenta becomes sensitized to infection late in gestation, when a population of GPCMV-susceptible cells develop near the base of the main placenta, and that the immune response to these infected cells compromises placental function.

In this study, we used spatial transcriptomics to compare gene expression in age-matched GPCMV-infected and control placentas at a near-cellular resolution. We found that infection affects gene expression in anatomic regions, such as the labyrinth, that are not directly infected by the virus. When gene expression was compared between GPCMV-infected cells and their immediate microenvironment and a comparable area in mock-infected placentas, we found that the local inflammatory response to GPCMV was driven by upregulated chemokine gene expression. Similarities in the placental response to CMV infection between humans and guinea pigs indicate that the rodent could be a valuable model for testing immunomodulatory therapies that aim to prevent fetal injury after chronic placental infections.

## Results

### Spatial transcriptomics of the guinea pig placenta

HCMV and GPCMV both cause focal infections in the placenta [10, 33, 38]. When virus-specific immunohistochemistry or *in situ* hybridization is used to detect GPCMV, individual infected cells or small groups of infected cells can be detected in both the placenta and decidua but are rare [33, 38–40]. However, we have found that comparably large areas of infected cells can be consistently detected at the base of the main placenta when dams are infected after mid-gestation and euthanized near term [38]. These cells are often found adjacent to the subplacenta or maternal arterial channels that perfuse the placenta. We utilized the Visium Spatial Gene Expression platform (10x Genomics) to study transcription in the guinea pig placenta and decidua at a near single-cell resolution and illuminate how infection directly and indirectly affects gene expression. This next generation sequencing-based technology uses glass slides tiled with 55 µm diameter, oligonucleotide-containing spots; each capture area contains 5000 of these spots in a 6.5 x 6.5 mm^2^ square. The oligonucleotides include location-based barcodes, unique molecular identifiers, and poly-T sequences that are used to capture spatial transcriptomes [41]. **Figure 1** summarizes the experimental approach used in this project. Placentas were collected from guinea pigs that were bred during postpartum estrus, either GPCMV- or mock-infected at 35 days gestation (dGA), and euthanized 21 days post-infection (56 dGA) [42]. One half of each placenta was flash frozen in OCT compound for histopathologic studies and spatial transcriptomics (ST). A portion of the remaining placenta was collected for DNA extraction; extracted DNA was used for viral load quantification and fetal sex screening by droplet digital PCR (ddPCR) and endpoint PCR [43, 44]. For ST, the flash frozen tissue was sectioned onto Visium slides, hematoxylin and eosin (H&E) stained, and imaged. The tissue was then permeabilized and an Illumina sequencing library was generated from cDNA synthesized using the oligonucleotides that are tiled on the capture area.

**Figure 1.**
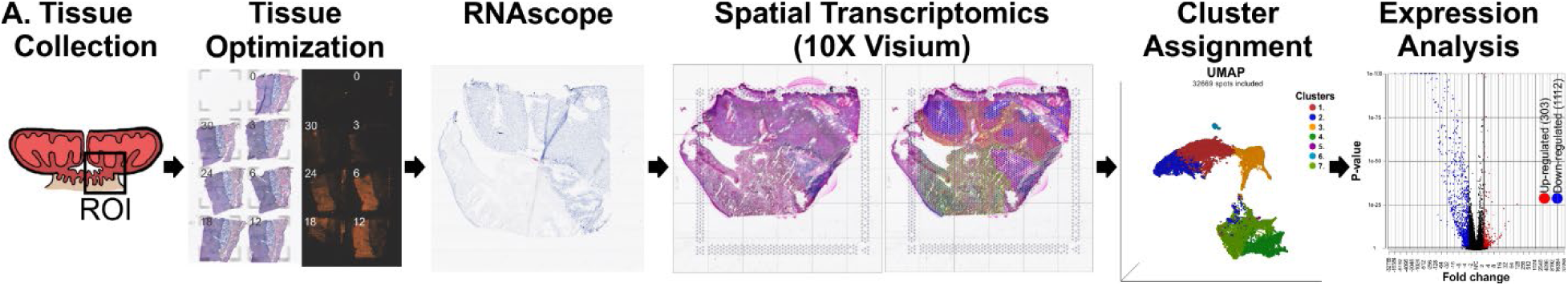
Spatial transcriptomics workflow. Gestational age-matched placentas from GPCMV- and mock-infected guinea pigs were divided and flash frozen in OCT compound. Visium Tissue Optimization Slides were used to determine the ideal incubation time for tissue permeabilization and on-slide cDNA synthesis. GPCMV-specific RNAscope was used to screen placentas for GPCMV-infected cells and select representative tissue to analyze on Visium Gene Expression Slides. Placentas were sectioned onto these slides, H&E stained, and imaged. After on-slide cDNA synthesis and Illumina sequencing, data was analyzed using Space Ranger and Partek Flow software. Gene expression in GPCMV- and mock-infected tissue was compared after graph based clustering and spatial transcriptomes that contained infected cells were identified based on their viral read content.

On-slide cDNA synthesis was optimized using Visium Tissue Optimization Slides (10x Genomics). Fluorescence microscopy revealed that more cDNA was produced from the main placenta than either the decidua or the subplacenta, regardless of how long the tissue had been incubated with permeabilization buffer (**Fig. 1**). This reflects the higher RNA content of the placenta, and we have observed that an average of 2.5±0.71-fold more RNA is extracted per mg of placenta than decidua in other unpublished experiments.

RNAscope specific to the viral transcript *gp3* was used to screen for infected cells and identify representative GPCMV-infected samples (**S1 Fig.**); five placentas (three female and two male) collected from three dams were selected for ST. GPCMV-infected cells were detected at the base of the main placenta in all these placentas. Six age-matched placentas from three mock-infected dams were used as controls. The guinea pig placenta is significantly larger than 6.5×6.5 mm^2^ at 56 dGA and each placenta was trimmed to so that portions of the placenta, decidua, and subplacenta would be included on each capture area. However, examination of the H&E-stained gene expression slides revealed that the subplacenta was not present in several tissue sections (**S2 Fig.**). To verify that infected cells were present in each capture area, a serial section was collected from each trimmed placenta and used for a second round of RNAscope (**S3 Fig.**). In the initial RNAscope analysis, one placenta (2395B) had a large area of *gp3^+^* cells in the decidua; this placenta was divided and sectioned onto two capture areas. Sample 2395B1, which should have corresponded to this area of infected decidua, did not contain gp3^+^ cells in the second round of RNAscope. GPCMV infected cells were detected where expected in all other samples.

cDNA from the twelve capture areas was used to prepare sequencing libraries, pooled, and sequenced. Apart from sample 735B, more than 5X10^4^ reads were sequenced per spot and a median of 3900 host transcripts were detected in each spatial transcriptome (**S1 Table**). In GPCMV-infected placentas, a median of 1.45X10^5^ (range 3X10^3^ to 2.6X10^5^) reads mapped to the viral genome per sample. These reads mapped to the 3’ end of viral transcripts and spanned the entire GPCMV genome (**Fig. 2**). Fewer than 100 reads mapped to the viral genome in mock-infected tissue sections and these reads were generally highly repetitive sequences.

**Figure 2.**
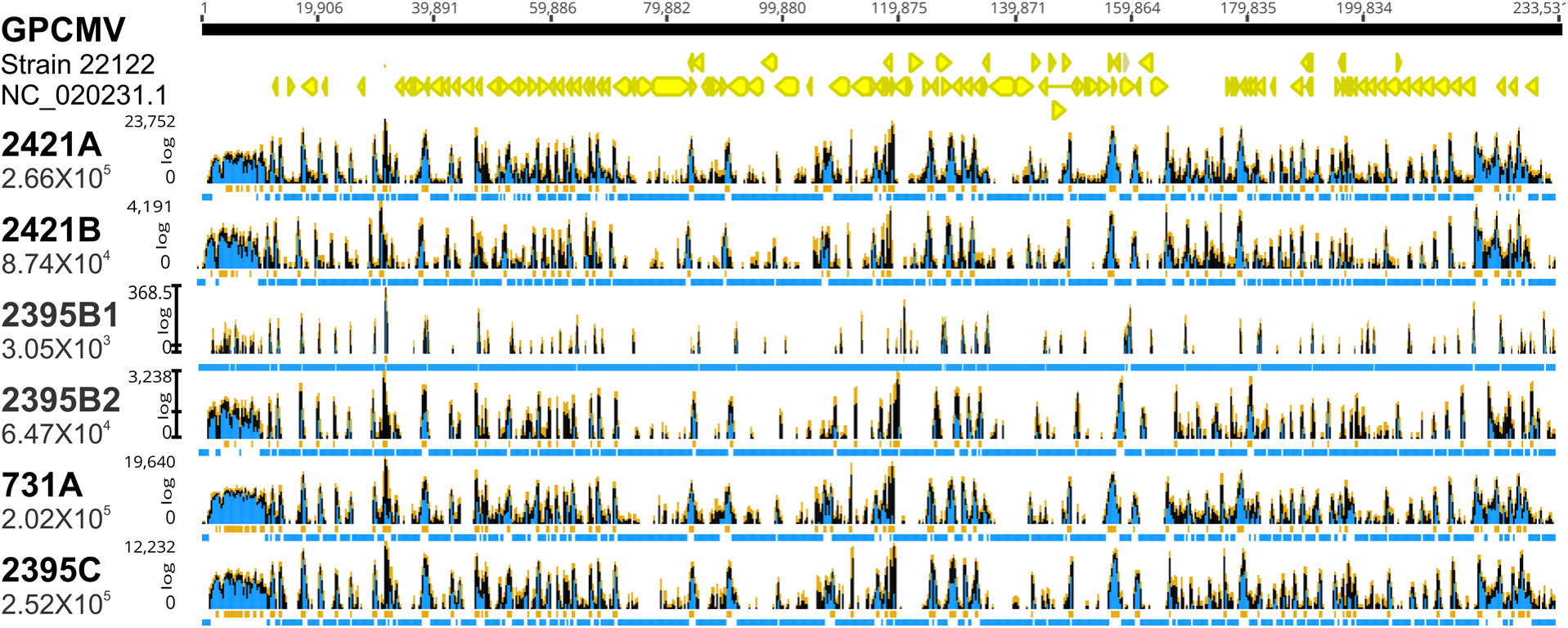
GPCMV sequences detected in spatial transcriptomes aligned to the viral genome. Sequencing reads were aligned to the GPCMV genome (NC_020231.1) using SpaceRanger and visualized with Geneious (v11.1.5). The total number of reads mapping to the viral genome in each sample is shown. Coverage at each nucleotide position is graphed in log scale. Regions with coverage <1 read are highlighted in blue and >100 in orange.

### GPCMV infection alters gene expression across the distinct anatomic regions of the guinea pig maternal-fetal interface

A combination of graph-based clustering and manual refinement was used to compare gene expression across the different anatomic regions of the guinea pig placenta. The 3000 most variant guinea pig transcripts were used for graph-based clustering; viral transcripts were excluded from this analysis. Ten clusters were produced when a resolution setting of 0.5 was used for graph-based clustering (**S4 Fig.**). Cluster assignment generally reflected where spatial transcriptomes originated at the maternal-fetal interface. The computationally generated clusters were then manually refined. Firstly, data from spatial transcriptomes located in folded tissue or near large blood spaces were excluded from later analyses. Three clusters that included the interlobar or marginal syncytium were combined into a single cluster representative of all non-capillarized syncytium and three clusters were combined and classified as parts of the placental stem and junctional zone. Seven clusters remained after manual spot reassignment (**Fig. 3**). While this underrepresents the complexity of the placenta, a conservative approach allowed us to compare gene expression across many spatial transcriptomes per cluster. The capillarized syncytium was represented by two clusters that comprise the core and periphery of the labyrinth. The non-capillarized syncytium and subplacenta were each represented by a single cluster. The remaining transcriptomes, located in the junctional zone, the stem of the placenta, and the decidua, formed three clusters, the smallest of which was comprised of only 389 transcriptomes and was excluded from later analyses. Consistent with our observation that the subplacenta was not sampled in several tissue sections, spots assigned to the subplacenta cluster were not present in all twelve capture areas (**Fig. 4a**).

**Figure 3.**
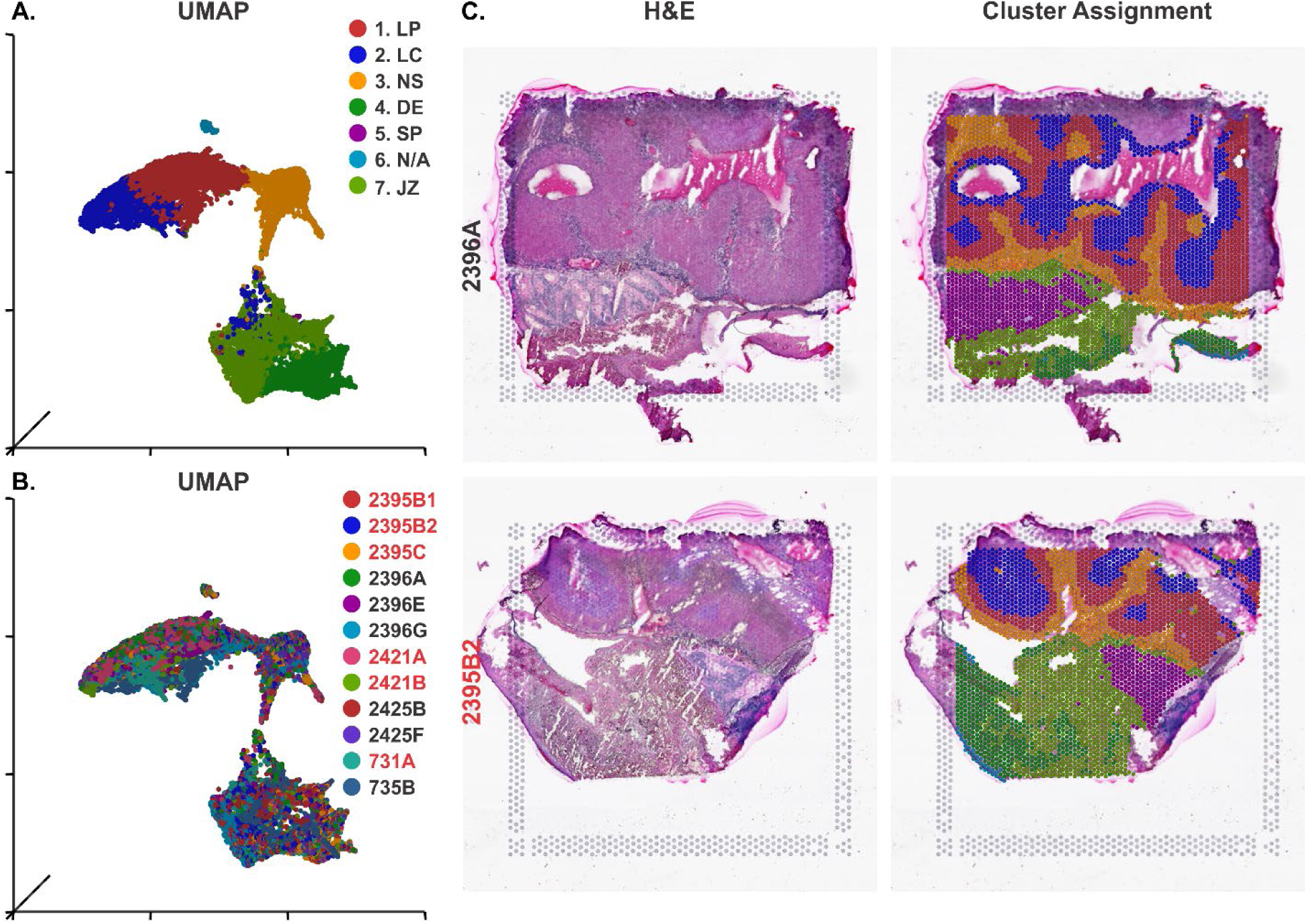
Gene expression in spatial transcriptomes reflects their location at the maternal-fetal interface. Spatial transcriptomes were classified using a combination of graph-based clustering and manual reassignment. (**A**) UMAP projection showing transcriptomes colored by the cluster assignment used for differential expression analyses. LP- labyrinth periphery, LC- labyrinth core, NS- non-capillarized syncytium, DE- decidua, SP- subplacenta, N/A- not assigned, JZ-junctional zone and placental stem. **(B)** UMAP projection showing spots colored by sample identifier, with GPCMV-infected samples labeled in red, illustrating the integration of data from three Visium slides and twelve capture areas. **(C)** Representative sections of mock- and GPCMV- infected placenta. The H&E stained tissue sections are shown on the left. Spatial transcriptomes were colored by cluster assignment in the panels on the right. Transcriptomes that were excluded from subsequent analyses are not colored.

**Figure 4.**
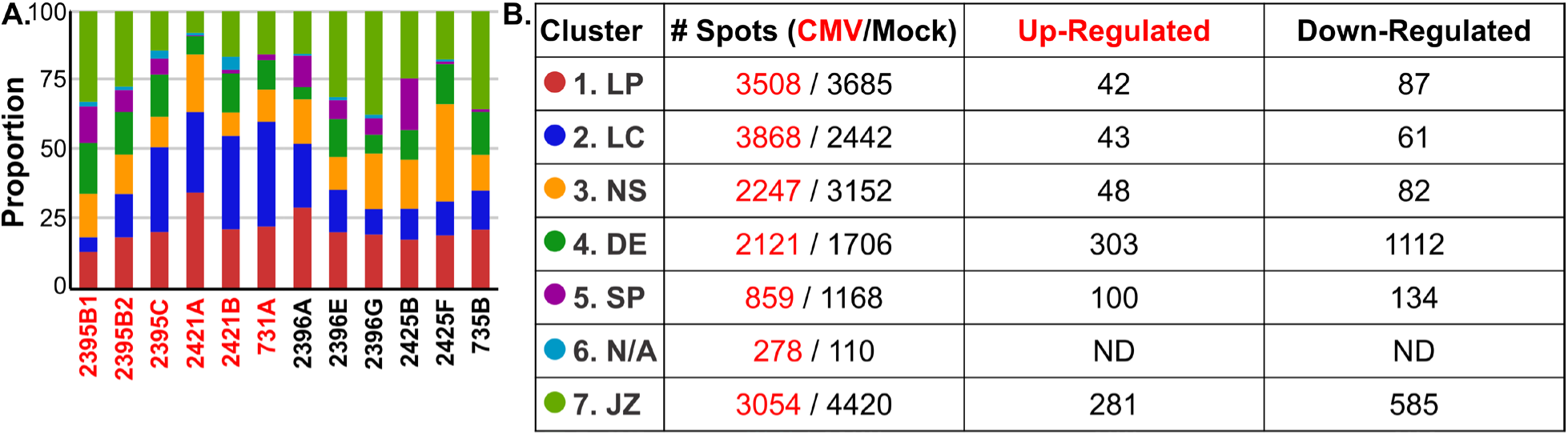
GPCMV infection dysregulates gene expression in the placenta and decidua. Gene expression was compared between sections of GPCMV- and mock-infected placenta (N=6/group). **(A)** The proportion of spatial transcriptomes in each capture area that were assigned to a given cluster is illustrated by bar chart. **(B)** Transcripts that were differentially regulated by GPCMV infection in each cluster were identified by gene-specific analysis (two-fold or greater change, *p* ≤0.05). The total number of spatial transcriptomes from GPCMV- and mock-infected capture areas included in this analysis and number of genes that were up- or down-regulated in each comparison are shown.

Cluster-specific biomarkers were identified by using Student’s t-tests to compare gene expression in the spatial transcriptomes assigned to one cluster with the combined data from all other spatial transcriptomes. **S2 Table** ranks the top 20 biomarkers for each cluster by significance and fold change. To illustrate how biomarker expression varies in the placenta and decidua, expression data for select biomarkers was overlaid onto images of representative control and GPCMV-infected placentas (**S5 Fig.**). The biomarkers for the two labyrinth clusters were partially overlapping. Conversely, the decidua and subplacenta clusters both had a high number of highly expressed biomarkers that were unique to the regions. Interestingly, several prolactin-like transcripts were identified as biomarkers for the labyrinth and non-capillarized syncytium. Guinea pigs, like murid rodents and ruminants, have an expanded prolactin locus and two of these transcripts, *Prlrp1* and *Prlrp2,* were previously shown to be expressed in the guinea pig placenta [45]. Our analysis found that *Prlrp1* was a biomarker for the labyrinth clusters and *Prlrp2* was enriched in the non-capillarized syncytium but expressed at a high level throughout the placenta and decidua. A third, previously uncharacterized prolactin-like transcript, *Ghc2* (LOC111754958), was a biomarker of the non-capillarized syncytium. These prolactin-like transcripts may have evolved to regulate placental development in guinea pigs.

To assess how GPCMV infection affected gene expression in the different regions of the placenta, the spatial transcriptomes of infected and mock-infected tissues were compared on a cluster-by-cluster basis. For this analysis, gene expression data was normalized to counts per million plus one and log_2_ transformed. Differentially regulated transcripts were identified by gene-specific analysis. Broadly, GPCMV infection downregulated more host transcripts than it upregulated (**Fig. 4b**). Functional enrichment analyses of the differentially regulated transcripts was done with g:Profiler [46]. Infection had a comparably modest effect on gene expression in the labyrinth, the non-capillarized syncytium, and the subplacenta. Several terms related to an antiviral response, including immune response (GO:0006955), type I interferon production (GO:0032606), and chemokine activity (GO:0008009), were enriched in these three regions after GPCMV infection (**S5 Fig.**). Terms related to metabolic functions such as peptidase activity, sterol transport, and signaling receptor binding and activity also were enriched in these clusters. GPCMV-infected cells were rarely detected in these regions by RNAscope and few of the spatial transcriptomes assigned to these clusters contained viral reads. Thus, it can be inferred that infection-associated changes in gene expression in the labyrinth, non-capillarized syncytium, and subplacenta are largely indirect effects of viral infection elsewhere in the placenta.

GPCMV infection affected markedly more transcripts in the two clusters that mapped to the decidua or placental stem and junctional zone. Most GPCMV-infected cells were assigned to the later cluster and 74 immune response (GO:0006955) transcripts, including many involved in chemokine signaling, were dysregulated by infection (**Fig. S5**). Viral infection downregulated transcripts with functions related to protein binding (GO:0005515), lipid transport activity (GO:0005319), small molecule metabolic processes (GO:0044281), and developmental processes (GO:0032502) in these tissue compartments. Taken together, this differential gene expression analyses revealed how GPCMV infection affects regions of the placenta that are not directly infected. While GPCMV infection dysregulates fewer transcripts in the main placenta than in the decidua, placental stem, or junctional zone, the upregulation of *Irf7* and select interferon stimulated genes (e.g. *Isg15*, *OasL*, and *Rsad2*) provides evidence that an antiviral response occurs throughout the placenta. The dysregulation of transcripts involved in metabolic or developmental processes suggests that GPCMV infection reprograms gene expression throughout the decidua and placenta in ways that could be harmful to fetal development.

### Gene expression changes in the GPCMV-infected microenvironment

GPCMV transcripts are polyadenylated and spatial transcriptomes that contained viral reads were identified in all six samples of GPCMV-infected placenta (**Fig. 5**). To determine how GPCMV affects transcription in infected cells and cells in their immediate vicinity, we compared the results of *gp3*-specific RNAscope with the number of GPCMV reads that were detected per transcriptome (**S2 and S6 Figs.**). Spots that contained more than one hundred viral reads (N=99) colocalized with *gp3+* cells in the serial tissue section and were annotated as GPCMV-infected. Sample 2395B1 was excluded from this analysis because it lacked spatial transcriptomes that contained more than one hundred viral reads and *gp3*-positive cells. Spots that were located within 110 µm of GPCMV-infected cells were classified as part of the GPCMV-infected microenvironment (N=285) for this differential gene expression analysis. In total, 384 spatial transcriptomes from GPCMV-infected placentas were compared with 1148 spatial transcriptomes from the mock-infected tissue sections that were similarly located in the placenta **(Fig. S6)**. Most spatial transcriptomes included in this analysis had been assigned to either the placental stem and junctional zone or the non-capillarized syncytium clusters (**Fig. 6a**).

**Figure 5.**
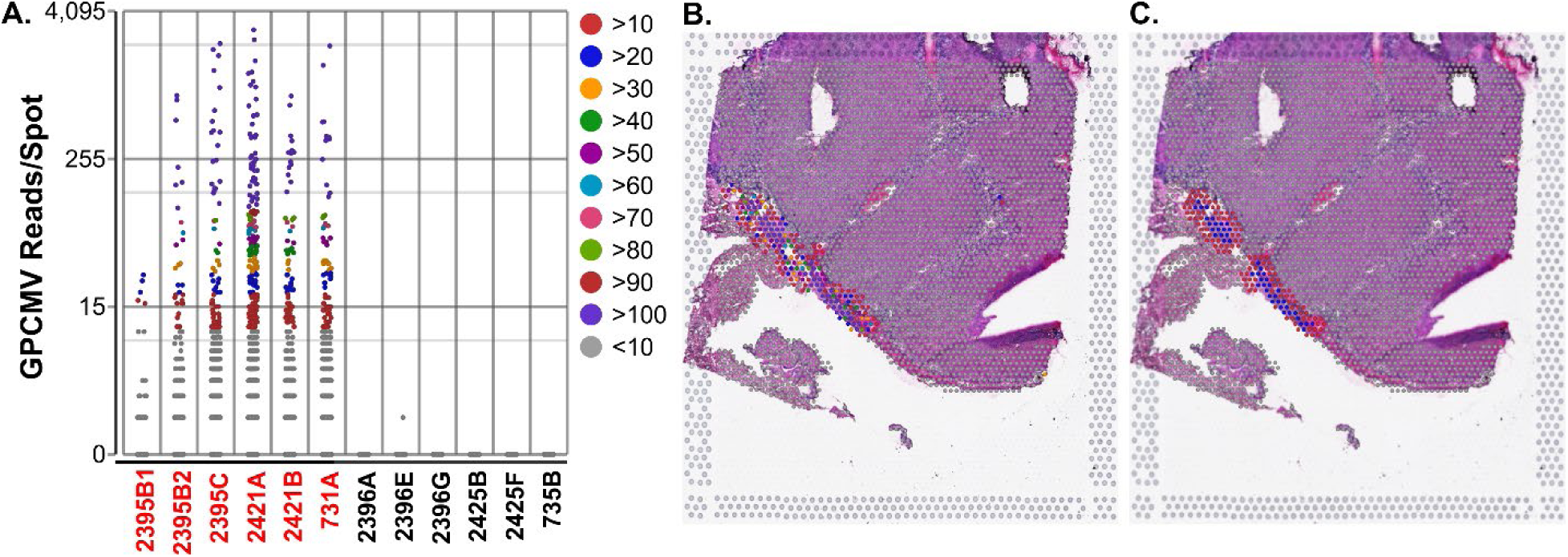
GPCMV-infected cells are detected in spatial transcriptomes. SpaceRanger was used to map reads to the GPCMV genome and spatial transcriptomes that contained viral transcripts were identified. (**A**) The number of GPCMV-specific reads in each transcriptome was plotted by sample identifier. GPCMV-infected samples are noted in red, and the data was colored by the viral read count. (**B**) Sequencing data was overlaid onto an image of a GPCMV-infected sample (2421A). Spots were colored based on GPCMV-read count as in (A). (**C**) For differential gene expression analyses, transcriptomes that contained >100 GPCMV reads were defined as “infected” (blue) and any spot within 100 µM was classified as in the GPCMV-infected microenvironment (red).

**Figure 6.**
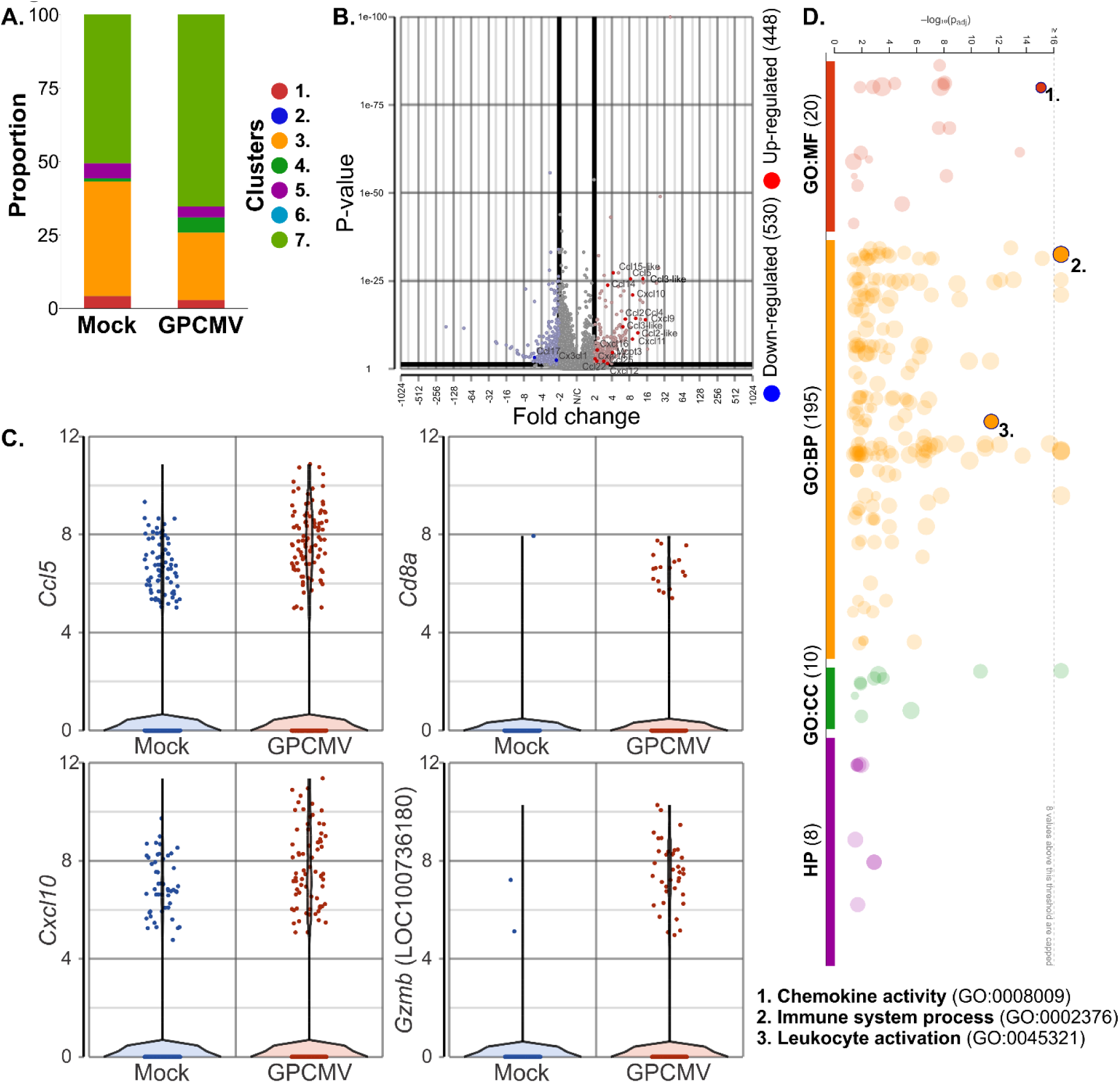
Chemokine and T cell-associated transcripts are upregulated in or near GPCMV-infected cells. Gene expression was compared between spatial transcriptomes that contained ≥100 GPCMV reads or were within 110 µm of infected cells and a comparable area in mock-infected tissue. **(A)** Bar graph illustrating the cluster assignments for the two groups of transcriptomes that were compared in this analysis. (**B**) Transcript abundance illustrated by volcano plot, highlighting transcripts with chemokine activity (GO:0008009). (**C**) Violin plots displaying the expression of select chemokine transcripts or transcripts related to a cytotoxic immune response in individual transcriptomes. (**D**) g:Profiler was used for functional enrichment analysis to identify terms that are enriched among the transcripts that are differentially regulated by GPCMV infection. Select gene ontology terms are highlighted on a modified Manhattan plot.

448 guinea pig transcripts were upregulated in or near GPCMV-infected cells and 530 were downregulated (≥2-fold change, *p*≤0.05, **Fig. 6b**). Gene ontology analysis revealed a transcriptional signature indicative of immune activation in the GPCMV-infected tissues (**Fig. 6c**). Notably, the inflammatory response to GPCMV infection appears to be predominantly driven by chemokine signaling. 16 of 18 chemokine transcripts that were enriched in or near GPCMV infected cells were upregulated, including multiple chemokines that signal through Cxcr3 and several chemokines (*Ccl2*-like [LOC100717552], *Ccl3*-like [LOC100734903 and LOC100735187], and *Ccl15*-like [LOC101788094]) that may be unique to guinea pigs (**Fig. 6b**). 67 transcripts involved in leukocyte activation (GO:0045321), including markers of a cytotoxic immune response such as *Cd3d*, *Cd8a*, and granzyme transcripts, were also more highly expressed at sites of GPCMV infection. Cytokines that are associated with acute placental infection (e.g. IL-1β, TNFα, IL-6, and IL-8) were not differentially regulated by GPCMV. Similarly, while numerous interferon stimulated gene transcripts (e.g. *Isg15*) were upregulated by infection, transcripts encoding interferons were not affected by GPCMV infection. Cumulatively, spatial transcriptomics resolved direct and indirect effects of GPCMV infection in the placenta and has revealed the central role of chemokine signaling in the GPCMV-infected placenta.

## Discussion

Placental infection plays a central role in CMV pathogenesis during pregnancy. In this study, we used ST to investigate how GPCMV affects gene expression at the maternal-fetal interface. With its 55-µm spot resolution, the Visium Spatial Gene Expression Platform enabled the most detailed analysis of transcription in the guinea pig placenta and decidua to date, revealing how infection impacts transcription in infected cells and their immediate microenvironment and illuminating how regions of the placenta are affected by the virus when not directly infected.

Research in primary trophoblasts, tissue explants, and organoids has provided insights into how HCMV affects the maternal-fetal interface, but these models have notable limitations [18, 20, 25, 26, 47, 48]. During *in vitro* experiments, placental cells or tissue explants are generally exposed to high infectious doses of virus by non-physiologic routes. Samples are analyzed within days of the initial infection, and the resulting data may reflect features of the innate host response to HCMV. Bulk RNA seq experiments have found that HCMV elicits markedly different transcriptional responses from decidua and placenta [20, 26]. Moreover, first and third trimester trophoblasts and tissue explants are differentially susceptible and responsive to HCMV infection [5, 26, 49, 50]. Thus, how CMV infection can cause placental injury likely depends upon when infection occurs during gestation.

Clinically, HCMV causes chronic placental infections that are characterized by focal lesions and a heterogeneous population of maternal and fetal leukocytes respond to infection [4, 10]. This complexity cannot be easily modeled *in vitro*, but studying samples of infected placenta collected after congenital infections presents its own challenges. Unlike Zika or rubella viruses, primary HCMV infection does not present with diagnostic clinical signs (e.g. rash), molecular methods must be used to confirm infection during pregnancy, most cases are not identified until after birth. Sample heterogeneity further complicates the analysis of clinical samples. For example, ST was used to study placentas collected after SARS-Cov-2 infection during pregnancy [51]. Most of nine samples analyzed did not contain virus that could be detected by qRT-PCR, *in situ* hybridization, or ST. The highest viral loads and most significant infection-associated changes in gene expression were found in two placentas recovered after intrauterine fetal demise at significantly different times in pregnancy (22.3 versus 35.0 weeks). Experimental studies in animal models thus play an important role in revealing how CMV and other viral infections can affect the maternal-fetal interface.

Guinea pigs have long been used for preclinical CMV vaccine development, but the limited availability of species-specific reagents has been a barrier to more sophisticated studies of CMV pathogenesis [52]. Species-agnostic next generation sequencing technology has revolutionized discovery science in guinea pigs and other unconventional animal models. For example, scRNA-Seq revealed that naked mole-rats, a hystricomorph rodent like the guinea pig, lack conventional natural killer cells [53]. We have previously used bulk RNA-seq to examine how guinea pig amniotic epithelial cells respond to infection *in vitro* and assess how primary GPCMV infection during the embryonic period or after mid-gestation differentially affect the placenta [38, 43]. However, bulk RNA-seq and most other conventional gene expression analyses quantify transcript abundance in tissue homogenates. This effectively averages transcript abundance across numerous cell types and states. ST enables gene expression analysis at a near single-cell resolution while preserving cellular location in the tissue of interest [54]. While the fresh-frozen Visium Spatial Gene Expression workflow is not reliant on species specific reagents, the 55-µm resolution of each spatial transcriptome is a limitation of the technology. Depending on the cellular density in tissue being analyzed, each spatial transcriptome reflects the RNA content of one to ten cells. Computational approaches have been developed that can infer the cellular composition within spots, as demonstrated by recent studies that used sc- or snRNA-Seq datasets to study the human maternal-fetal interface [51, 55]. While cell atlases have been developed for the human and mouse placenta, a similar resource has yet to be developed for the guinea pig and an automated cell type classification was not done in this study. [56–59]

This study revealed that GPCMV infection dysregulates more transcripts in the decidua and junctional zone than in the labyrinth or non-capillarized syncytium. HCMV antigens and DNA are more frequently detected in decidua than in placenta, but GPCMV is only sporadically detected in the decidua by *in situ* hybridization at 21 dpi (**Fig. S1**) [38, 60]. Additional experiments are necessary to assess whether GPCMV infects the decidua at earlier times in infection and resolves. One observation from gene set enrichment analysis was that GPCMV infection downregulates lipid metabolism and transport in the maternal-fetal interface. Fetal development and growth are dependent upon the regulated transport of maternal nutrients, including fatty acids, across the placenta and maternal hyperlipidemia is typical during a healthy pregnancy [61, 62]. Abnormal maternal lipid levels are observed during pregnancy-induced hypertension, preeclampsia, and preterm birth [63–65]. At a cellular level, DNA viruses hijack lipid metabolism pathways to promote replication, and HCMV infection promotes fatty acid synthesis and modifies cholesterol synthesis and trafficking *in vitro* [66–70]. Given that ZIKV infection also perturbs lipid homeostasis in the placenta, altered lipid metabolism may a mechanism by which viruses cause placental insufficiency [71].

Placental dysfunction and adverse fetal outcomes can be caused by interferon signaling and other inflammatory responses to infection [8, 72]. Infection is a leading cause of preterm birth, and the acute inflammatory responses to ascending bacterial infection involves cytokines, including IL-1β, IL-6, IL-8, and TNF-α, that are normally upregulated during parturition [73]. Polyinosinic:polycytidylic acid (Poly I:C), a synthetic analog of the viral pathogen associated molecular pattern double-stranded RNA, can affect placental and fetal development in mice and rats depending upon the dose, route of administration, and the timing of treatment during pregnancy [reviewed in [74]]. Experiments in *Ifnar*-transgenic mice have demonstrated that fetal growth restriction and resorption can be caused by Type I interferon signaling that occurs in the Zika virus (ZIKV)-infected fetus and placenta [27, 75, 76]. In humans, Aicardi–Goutières (AGS) syndrome, a rare inherited disease characterized by dysregulated Type I interferon production, can present during pregnancy with ultrasound findings reminiscent of congenital infections [77].

Multiple studies have found that cCMV causes proinflammatory cytokines to accumulate in amniotic fluid, and elevated CXCL10 levels are a biomarker of congenital infection [22–24]. CMV infection also upregulates *CXCL10* transcription in tissue explants, amniotic epithelial cells, and decidual organoids *in vitro* [20, 26, 43]. Elevated amniotic fluid concentrations of CXCL10 also occurs during chronic chorioamnionitis and maternal anti-fetal rejection; the chemokine may drive T cell recruitment to the placenta and/or fetal membranes [78–80]. Bulk RNA-Seq found that *Cxcl10* is upregulated in the GPCMV-infected placenta, and ST confirmed this finding while also revealing that a broad mix of chemokines are upregulated in the microenvironment around GPCMV-infected cells [38]. Curiously, while individual interferon stimulated genes are upregulated during placental GPCMV infection, interferon transcripts were not regulated by infection. Interestingly, one study found no association between CXCL10 concentrations and the detection of HHV6, parvovirus B19, or Epstein-Barr virus in amniotic fluid [23]. Nonhuman primate studies also indicate that the inflammatory response to CMV may be distinct from other viral causes of placental and congenital infections such as ZIKV [81, 82].

Primates and rodents diverged 100 million years ago, and the cytomegaloviruses have been cospeciating with their hosts ever since [83, 84]. The species-specificity of HCMV precludes the direct study of the virus in animal models, and while *in vivo* experiments using closely related primate and guinea pig cytomegaloviruses can reveal how infection affects the placenta under controlled experimental conditions, virus and host-specific immune adaptations could limit the translatability of these studies to humans. For example, GPCMV-specific adaptations to its host’s immune system include the convergent evolution of class I MHC mimics and a unique viral chemokine, *gp1* [85, 86]. Guinea pigs and other caviomorph rodents have a unique population of mononuclear cells, Kurloff cells, with natural killer (NK) cell activities [87]. While direct functional comparisons to human or mouse decidual NK cells have yet to be completed, the abundance of circulating Kurloff cells is regulated by estrogen and the cells localize to the placenta and uterus in pregnant animals [88, 89]. While this study and our previous bulk RNA-seq of the guinea pig placenta provide valuable insights into the biology of the maternal-fetal interface in guinea pigs, more extensive studies of placental gene expression and guinea pig immunity are merited to support the use of the rodent as a comparative model for human gestation [38].

## Materials and Methods

### Ethics Statement

All animal procedures were conducted in accordance with protocols approved by the Institutional Animal Care and Use Committee (IACUC) at the University of Minnesota, Minneapolis (Protocol IDs: 1810-36403A and 2106-39180A). Experimental protocols and endpoints were developed in strict accordance to the National Institutes of Health Office of Laboratory Animal Welfare (Animal Welfare Assurance #A3456-01), Public Health Service Policy on Humane Care and Use of Laboratory Animals, and United States Department of Agriculture Animal Welfare Act guidelines and regulations (USDA Registration # 41-R-0005) with the oversight and approval of the IACUC. Guinea pigs were housed in a facility maintained by the University of Minnesota Research Animal Resources, who were accredited through the Association for Assessment and Accreditation of Laboratory Animal Care, International (AAALAC). All procedures were conducted by trained personnel under the supervision of veterinary staff.

### Infection of Pregnant Guinea Pigs and Tissue Collection

GPCMV seronegative, pregnant guinea pigs were mock- or GPCMV- infected as previously described [38]. Briefly, two- to three-month-old strain 2 females were housed with strain 2 males until pregnancy could be confirmed by palpation. At this point, the strain 2 male was replaced with a strain 13 male, who remained with the female until three days postpartum to establish a second, timed pregnancy [42]. Progesterone ELISA and/or palpation was used to confirm the second pregnancy. At 35 days dGA, guinea pigs were injected subcutaneously into the scruff of the neck with 0.5 mL of GPCMV-containing PBS (2×10^5^ PFU, SG13J2) or PBS alone [43]. Blood and plasma were collected from dams every seven days post-infection (dpi) until the animals were euthanized at 21 dpi. Several dams also received intraperitoneal (IP) injections of 1.5 mg of rat IgG (2 mg/mL in PBS) at days 0, 7, and 14 post-infection or an IP injection of 80 mg of Hypoxyprobe 1 (pimonidazole HCl) dissolved in PBS that was administered 90 minutes before the animal was euthanized (Supplementary Table 1). During tissue collection, the uterus was dissected, the decidua gently separated from the uterus, and each placenta was halved. One half of each placenta was embedded in optimal cutting temperature (OCT) compound and flash frozen in isopentane chilled with a dry ice ethanol bath. A portion of the remaining tissue was frozen at −80°C for DNA extraction and viral load quantification.

### Screening Placentas for GPCMV Infection and Fetal Sex

DNA was extracted from placentas using the DNeasy Blood and Tissue Kit (Qiagen). The abundance of GPCMV genomes in the tissue was quantified by ddPCR using primers and probes specific to *GP54* and the Bio-Rad QX200 system as previously described [43]. A multiplexed endpoint PCR assay that targets *Sry* (Y-specific) and *Dmd* (X-specific) was used to determine the sex of each placenta [44]. The location of GPCMV-infected cells in placenta was determined using an RNAscope^®^ assay specific to the viral transcript *gp3* (Probe V-CavHV-2-gp3) [38]. Fresh-frozen tissue was prepared for single-plex RNAscope® 2.5 assays as described in technical note 320536-TN Date 11112016. 10-μm sections of placenta were mounted onto Superfrost Plus slides (ThermoFisher), fixed using 4% paraformaldehyde, and dehydrated by sequential immersion in 50%, 70%, and 100% ethanol. After the slides were dried, the tissue was treated with RNAscope® hydrogen peroxide and Protease IV. The slides were then stained with RNAscope® 2.5 HD Detection Reagent – RED (322360-USM Rev. Date 11052015) as previously described (9).

### Spatial Transcriptomics

To optimize on-slide cDNA synthesis for ST, mock-infected placentas collected at 42 and 56 dGA were analyzed using Tissue Optimization Slides and Reagents (10x Genomics, PN-1000194). Blocks of frozen placenta were trimmed with the goal of including the main placenta, placental stem, and decidua on each capture area. 10 μm tissue sections were placed onto seven of the eight capture areas on the tissue optimization slide (CG000240 Rev B). Tissue was fixed with methanol, H&E stained, and imaged with a Huron TissueScope LE (CG000160 Rev A). For tissue permeabilization and fluorescent cDNA synthesis, capture areas were incubated with Permeabilization Enzyme for times that ranged from 3 to 30 minutes (CG000238 Rev C). After reverse transcription, slides were imaged with a Nikon Eclipse fluorescent microscope. An 18-minute permeabilization time was selected for ST experiments based on the quality of the fluorescent signal. Irrespective of the permeabilization time, more signal was observed in the main placenta than in the subplacenta, placental stem, or decidua.

Five GPCMV-infected and six control placentas were sectioned onto the twelve capture areas of three Visium Spatial Gene Expression Slides. These tissues originated from three GPCMV- and three mock-infected dams and were a mixture of males and females. Blocks were roughly trimmed to 6.5×6.5 mm^2^ so that a similar region of interest would be included in each capture area and, for GPCMV-infected samples, infected cells would be included in the capture areas. One GPCMV-infected placenta (2395B) was divided and sectioned onto two capture areas. The slides were fixed, H&E stained, and imaged with a Huron TissueScope LE. The tissue was then permeabilized for 18 minutes and cDNA synthesis, amplification, and sequencing library preparation was completed using the manufacturer recommended protocol (CG000239 Rev C). Visium libraries were combined and sequenced on a single Illumina NovaSeq S4 lane (2 x 150 PE run format). Sequencing results are summarized in Supplementary Table 1. A median of 2.2X10^8^ reads per capture area (range 1.0X10^8^ to 6.9X10^8^) and 7.3X10^4^ reads per spatial transcriptome (range 3.3X10^4^ to 1.8X10^5^) were sequenced. Sequencing data has been uploaded to the Gene Expression Omnibus (Series GSE303764, https://www.ncbi.nlm.nih.gov/geo/query/acc.cgi?acc=GSE303764).

### Data analysis

To analyze ST data, a custom reference sequence that combined the guinea pig genome (Cavpor3.0, GCF_000151735.1, Annotation Release 103), the GPCMV genome (strain 22122, NC_020231), and a list of Y chromosome transcripts, was prepared [90, 91]. Space Ranger (10x Genomics, version 1.3.0) was used to process the raw Illumina sequencing data. Space Ranger outputs were uploaded to *Partek*® *Flow*® software (v10.0). Reads that aligned to the GPCMV genome from each sample of infected placenta were visualized with Geneious (v11.1.5). Each capture area was manually examined and 927 spatial transcriptomes that were located under folded or visibly damaged tissue were excluded from downstream analyses. For graph-based clustering, data from the remaining 33,687 spots was filtered to the 3000 most variable features, normalized with SCTransform, and batch corrected with Seurat3 integration [92, 93]. 50 principal components were used for graph-based clustering and data presentation by Uniform Manifold Approximation and Projection (UMAP) dimensional reduction. Sensitivities from 0.3 to 1.0 were evaluated using the Louvain clustering algorithm. Higher resolution settings predictably identified more clusters but also resulted in considerable variation in the number of spatial transcriptomes assigned to each cluster across the twelve samples. A resolution setting of 0.5 was selected for subsequent analyses. The data was further refined by excluding 1018 spatial transcriptomes that were located at the edges of tissue sections and combining several clusters. Cluster-specific biomarkers were identified using Partek Flow’s “compute biomarker” tool. This algorithm compared the expression of features in each cluster with the expression observed in all other clusters combined using student’s t-tests.

To identify host transcripts that were differentially regulated by GPCMV infection, the abundance of all transcripts that were detected in >0.1% of spatial transcriptomes was normalized to counts per million plus one and log_2_ transformed. Differentially regulated transcripts were identified by gene-specific analysis (GSA). In one analysis, transcripts that were differentially regulated by infection were identified in each of the previously identified clusters. In a second analysis, spatial transcriptomes that contained ≥100 GPCMV transcripts and every spatial transcriptome within 110 µm were defined as the GPCMV-infected microenvironment. Transcript abundance was compared with similar area that was manually selected from mock-infected tissue sections, and 384 spatial transcriptomes from GPCMV-infected placentas were compared with 1148 spatial transcriptomes from control tissue. For both workflows, functional enrichment analyses on differentially regulated transcripts (twofold or more, *p* < 0.05) were completed using g:Profiler [46].

## Acknowledgements

This work was supported by the National Institutes of Health’s National Center for Advancing Translational Sciences, which provided funding for the Biorepository and Laboratory Services Team, and the resources and staff at the University of Minnesota University Imaging Centers (SCR_020997) and the Genomics Center (https://genomics.umn.edu). The content of this article is solely the responsibility of the authors and does not necessarily represent the official views of the National Institutes of Health’s National Center for Advancing Translational Sciences.

## Supporting Information

**S1 Figure.**
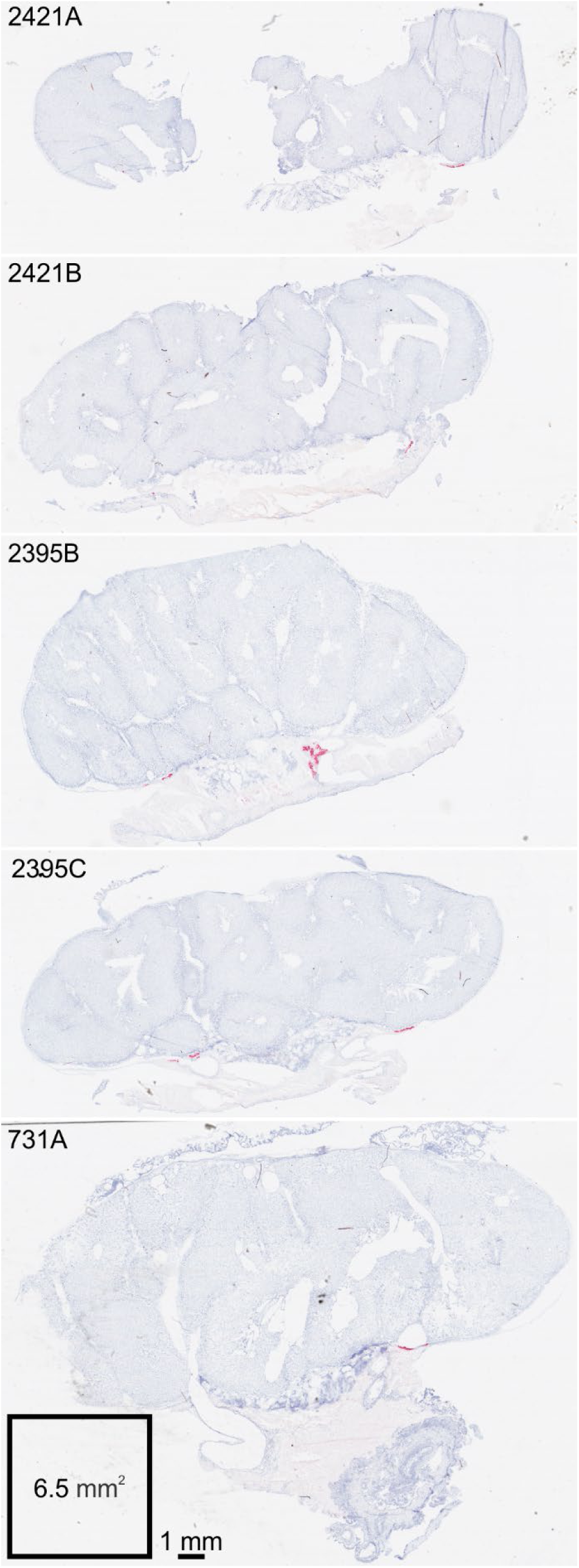
GPCMV-specific RNAscope. Fresh-frozen, GPCMV-infected placentas were cross-sectioned and stained with a RNAscope assay that detects the viral transcript *gp3* (red). Scans of the five infected placentas that were selected for ST are shown.

**S2 Figure.**
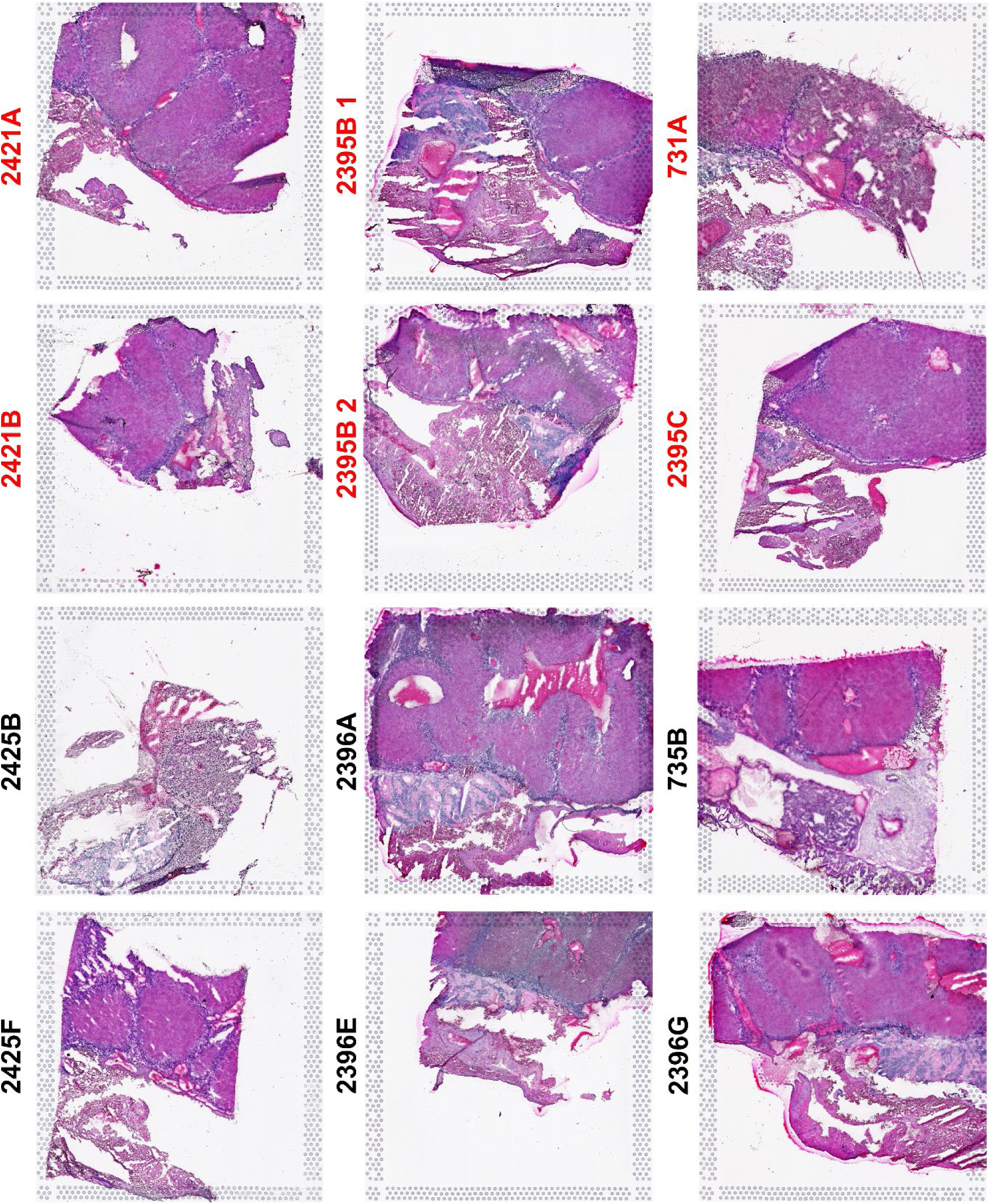
Hematoxylin and eosin-stained guinea pig placentas. GPCMV- (**Red**, N=5) and mock-infected placentas (N=6) were trimmed and sectioned onto twelve capture areas. The Visium Spatial Gene Expression Slides were H&E stained and imaged. The images were cropped to each capture area’s fiducial frame and oriented so that the main placenta is up and decidua down.

**S3 Figure.**
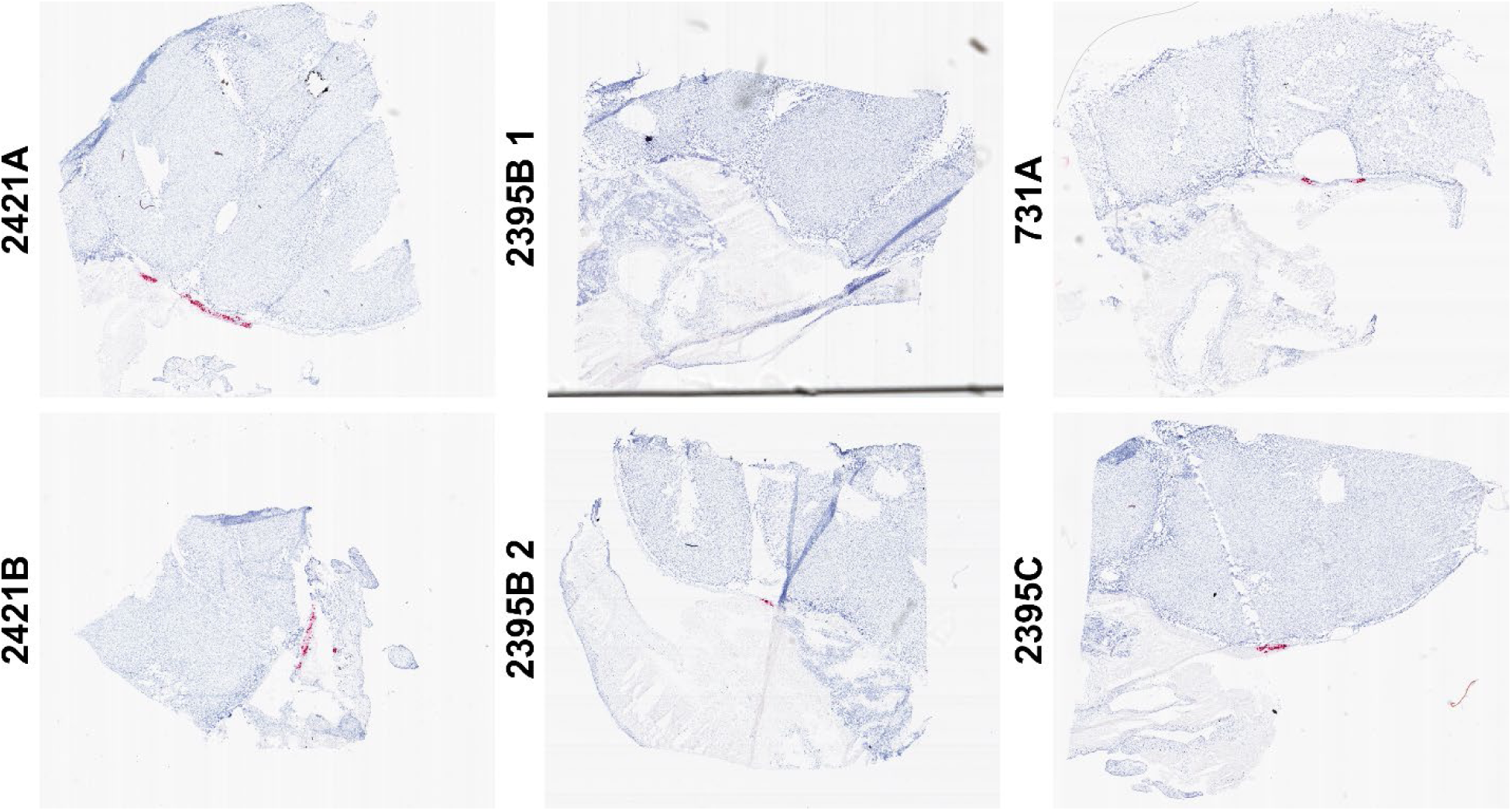
GPCMV-specific RNAscope of trimmed placentas. Placentas were trimmed to accommodate the 6.5X6.5 mm^2^ Visium capture area. After tissue was sectioned on the Visium Gene Expression Slides, the serial section from each sample was used for gp3-specific RNAscope (red) to confirm that infected cells had been placed within each capture area.

**S4 Figure.**
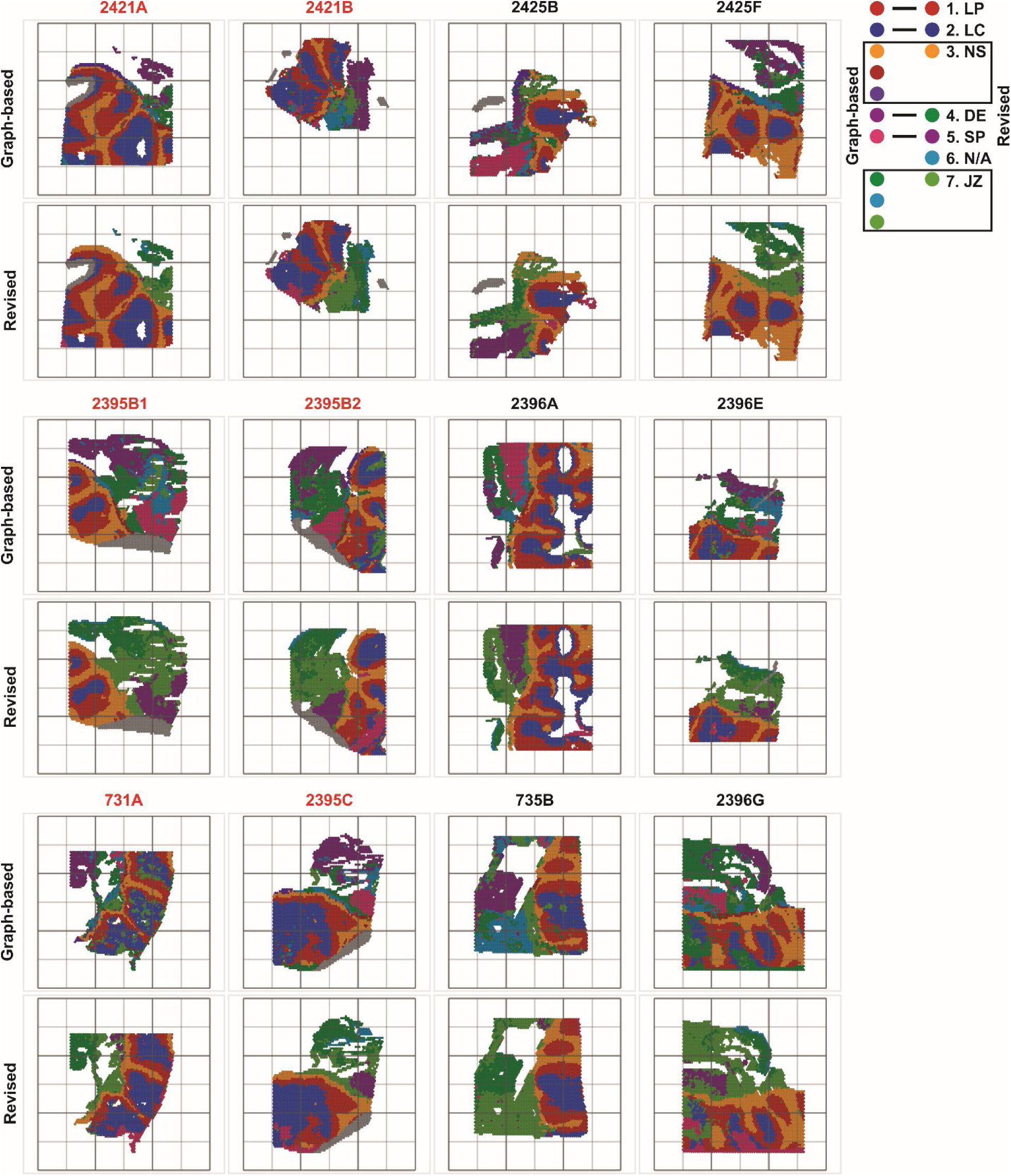
Spatial transcriptomes cluster by anatomic region. Graph-based clustering assigned spatial transcriptomes to ten groups. These were further manually refined to seven clusters, which included excluding data under folded tissue sections, near large blood spaces, and combining several smaller clusters into groups representing non-capillarized syncytium and the junctional zone and placenta stem.

**S5 Figure.**
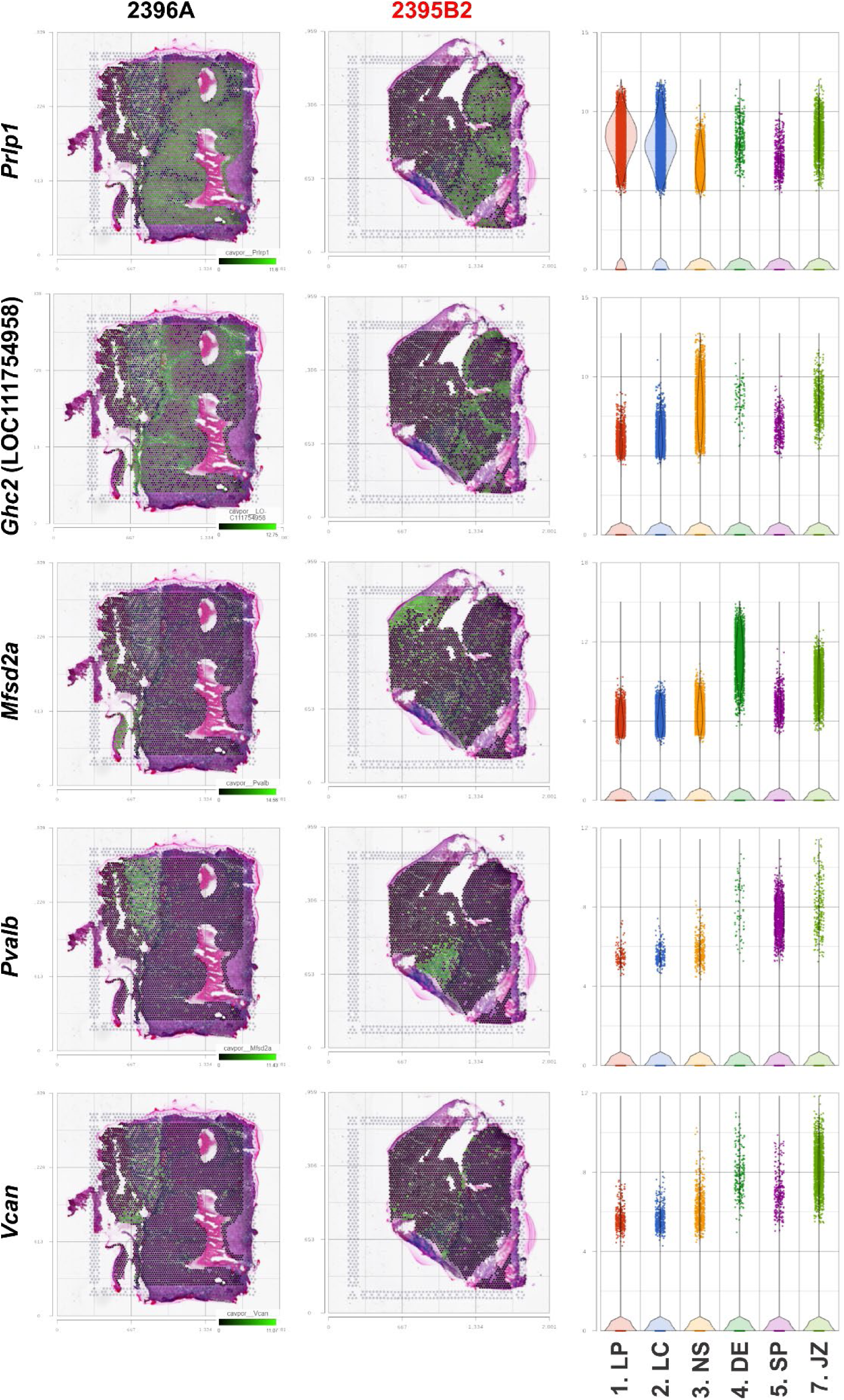
Biomarker expression in the anatomic regions of the guinea pig placenta. Transcripts were identified that were significantly enriched in each cluster. Gene expression data for select biomarkers was overlaid on two representative sections of placenta such that transcript abundance in each spot is represented by green intensity. Violin pots illustrate the expression of each biomarker in all samples with transcriptomes classified by cluster.

**S6 Figure.**
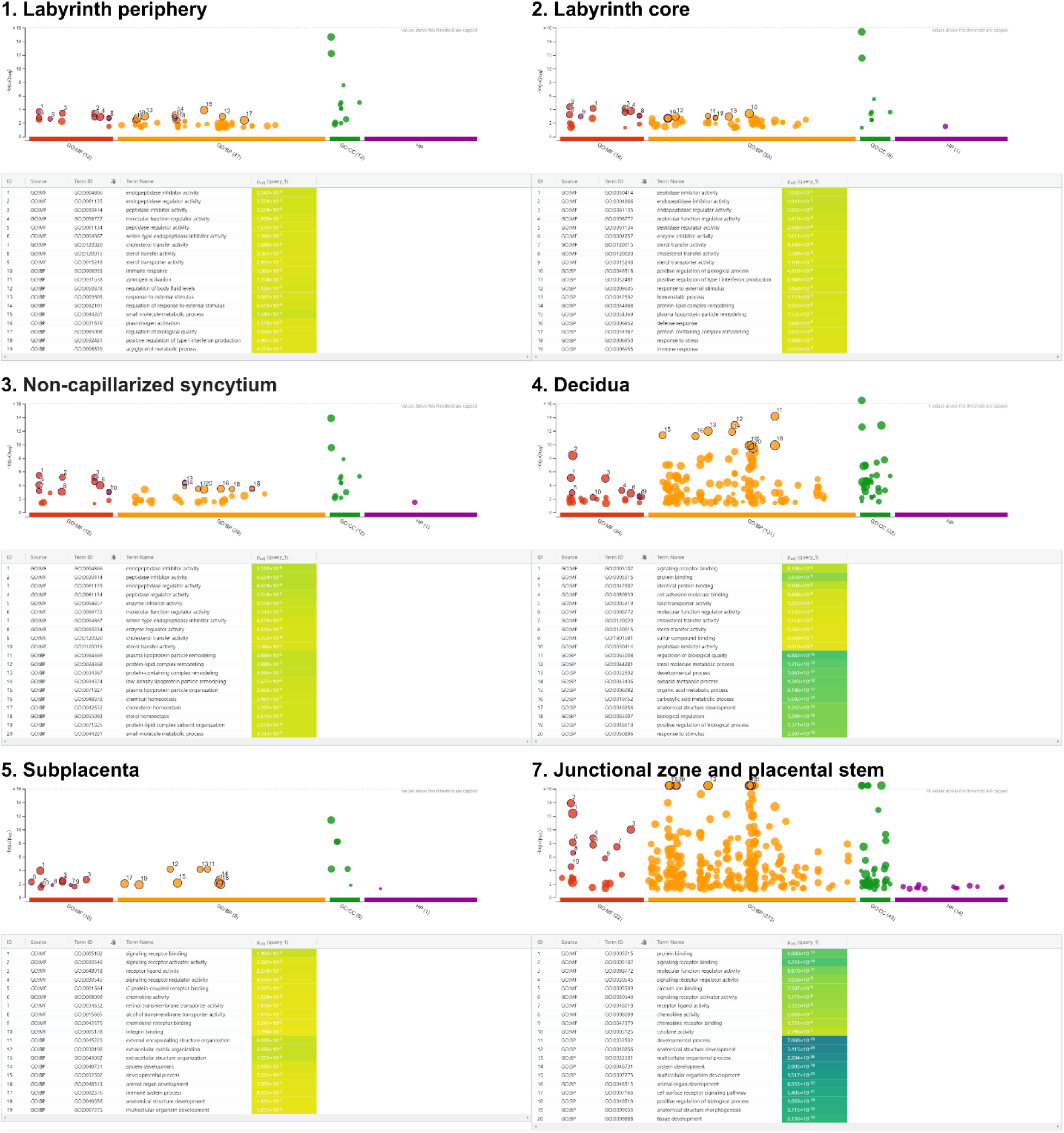
Functional enrichment analyses reveal pathways dysregulated by GPCMV infection. For each anatomic region of the placenta, transcripts that were significantly up- or down-regulated in GPCMV-infected placenta were identified using GSA. Functional enrichment analyses were done using g:Profiler [46]. Enriched pathways are illustrated by Manhattan plot and the ten most significantly enriched molecular functions and biological processes for each cluster are indicated.

**S7 Figure.**
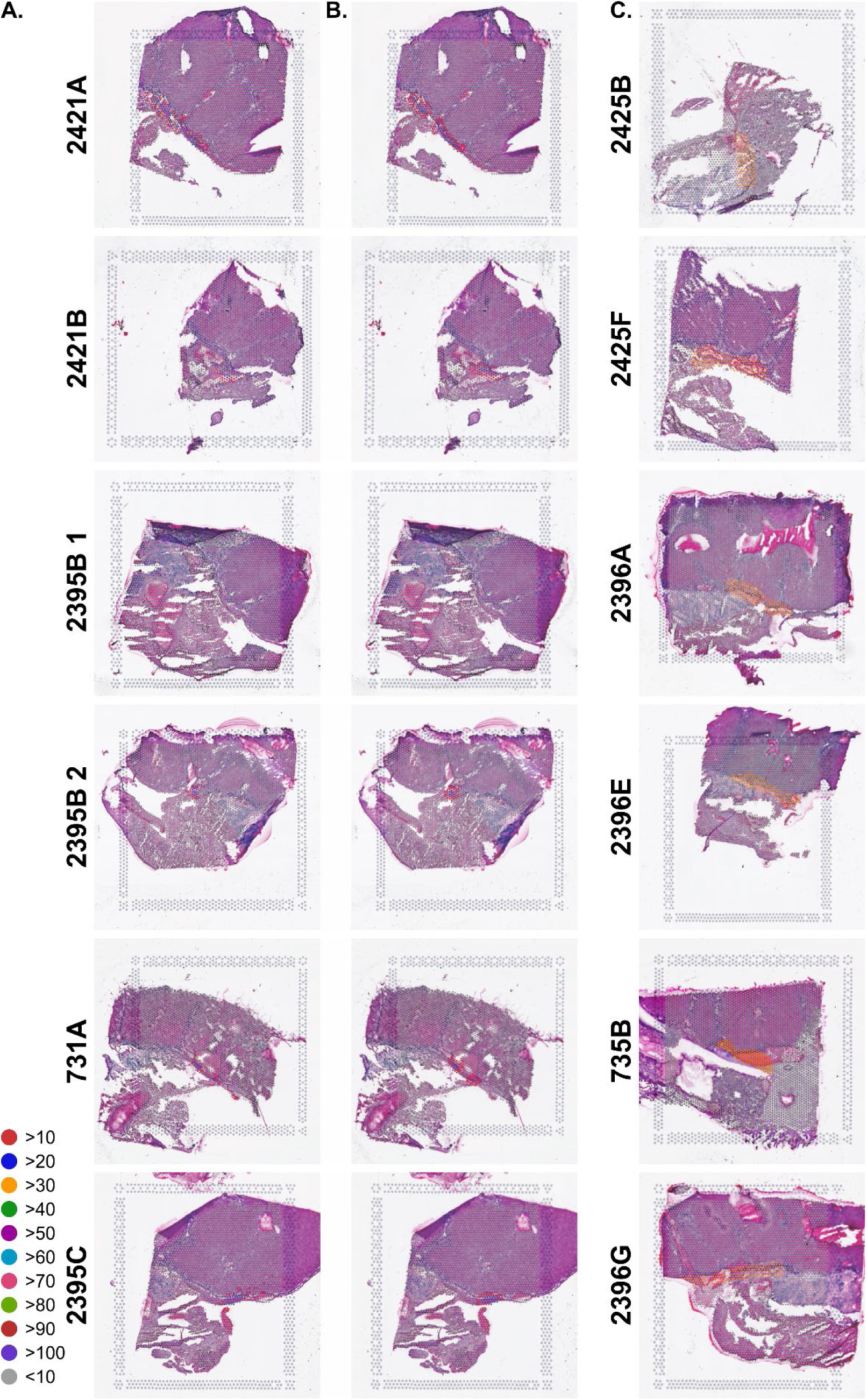
Identification of GPCMV-infected cells and their microenvironment in spatial transcriptomes. Transcriptomes that contained GPCMV reads were identified in each capture area. In GPCMV-infected samples, spatial transcriptomes were either colored by the abundance of viral reads or red for >100 reads or blue to designate the microenvironment near infected cells. In mock-infected samples, transcriptomes colored in yellow were selected for the differential gene expression analysis.

